# Probabilistic Approach to Understand Errors in Sequencing and its based Applications

**DOI:** 10.1101/2020.05.02.073650

**Authors:** Anuj Singh

## Abstract

Next Generation Sequencing has been applied in many areas of biology, including quantification of gene expression, Genome-Wide Association Study (GWAS), gene finding, Motif discovery and much more. Due to the vast area of application and importance in key findings in the field, massive genomic data is being generated using high throughput sequencing. Therefore, sequencing quality also needs to be evaluated in light of different applications. In order to develop effective diagnostic and therapeutic approaches, we need to accurately characterize and identify sequencing errors and distinguish these errors from their true genetic variant in sequencing, i.e. misreads follow a binomial distribution and it further can be approximated to the Poisson process for longer sequences. However, the insertion and deletion rates are 1000 times lower than substitution error rates and, therefore, less significant. The model assumes that error arrival at a position is not dependent on an error at other positions. Furthermore, errors in sequences can cause an error in studies based on multiple sequences and they also follow Binomial – Poisson Distribution (for example – Alignment is a merging of two Binomial processes for short sequences and it further can be approximated to Poisson for long sequences (for example – genomic sequence). It provides a systematic way to evaluate the accuracy in sequencing-based applications. Many error suppressing algorithms or techniques are there, and our Binomial Poisson model can provide a further systematic understanding of error behavior in short sequences so that more techniques for error removal can be developed with much efficient suppression rates.

## Introduction

For more than 30 years, most of biology’s research is based on DNA and its sequence, and therefore we have seen a rapid development in sequencing technologies. Because of higher degree parallelism and much smaller volumes, Next Generation Sequencing (NGS) achieves a much higher throughput with dramatically reduction in cost. This is the reason why the last decade has seen a steady increase of sequencing technologies in all fields of biology. Next Generation Sequencing has been applied in many areas of biology, including quantification of gene expression, Genome Wide Association Studies (GWAS), Gene finding, Motif discovery and much more. Studies such as PANCANCER (recently published in Feb’20) are totally based on large scale sequencing**[13]**. Due to the vast area of application and importance in key findings in the field, massive genomic data is being generated using high throughput sequencing. Therefore, sequencing quality also needs to be evaluated with respect to different applications. Apart from high throughput capability, the NGS has disadvantages of shorter reads and higher error rates compared to Sanger’s Sequencing and apart from cost, accuracy is also an important aspect.

By quality we are referring to errors in sequencing. By error, it is misread in a sequence. For example, there is a base *T* at a specific position but while sequencing it is read as *C*. So, there is an error at this position because it is misread while sequencing. The original sequence is different from the sequence we have and that is why it is called an error. There are three different possibilities for the base that could have been read as *G* or *A* or *C*. So any of the three cases would lead to error. In the same way, every base has three different possibilities of getting misread in the sequencing process and turns out to be an error. This particular type of error is termed as substitution error. For our paper, we are assuming that there is only substitution error in sequencing because insertion and deletion error rates are 1000 times lower than substitution rates**[3]**. Therefore, Substitution error rates are more significant than deletion and insertion. The substitution error rate by conventional NGS was reported to be >1% in 2011 and was found to be similar in later reports **[5].** Some groups tried to systematically investigate substitution error profile by analysing multiple datasets**[10].** Fox, Edward J et al also reported the average error rate to be 0.1% per nucleotide most of which is single nucleotide**[11]** Fields like comparative genomics use massive genomic data and any sequencing error may lead to a variation in results of the study. So it is very important to have a quantitative study of accuracy in a particular study. With this motivation, we first model the error in a single DNA sequence using a Binomial Process probabilistic model. Then we will approximate the poisson process for a genomic sequence. Further we will extend our model to find the probability of accuracy of the studies which include multiple sequences.

## Method

In this study, we are introducing a basic application of mathematical models for error analysis in DNA sequencing and its further applications. Many groups have previously applied probability in biology and worked towards new findings in the field. The hidden markov model for gene finding E Coli DNA is also an example of probability in biology**[1]**. It has played a major role in genetics and in this paper, we are introducing one more application to model the error in DNA sequencing.

DNA sequence is a sequence of *A, G, T* and *C*. The determination of this sequence is prone to errors and error can be led by various processes**[10,11]**. So while sequencing, occurence of error at a position is independent of occurrence of error at any other position. In simpler words, error at every position is independent. This would be our key assumption. For our model we are only considering substitution errors(as defined in Introduction). One more assumption that we are considering the probability of errors *A → G, A → C, A → T* and all types of substitution have the same probability of occurrence (because of average error rate)^#^.

Consider a DNA sequence of length ***n*** (as shown in the Figure 1). For every base in the sequence, we are defining two possible states as ‘***w****’* if the corresponding nucleotide is correctly sequenced and ‘***e***’ when there is an error in sequencing for the respective base. As a result, we have a state sequence of the same length (Figure 1) for our given DNA sequence where 5th and the ith position are in error state whereas the rest of the sequence is in correct state(correctly sequenced).

**Figure 1:**
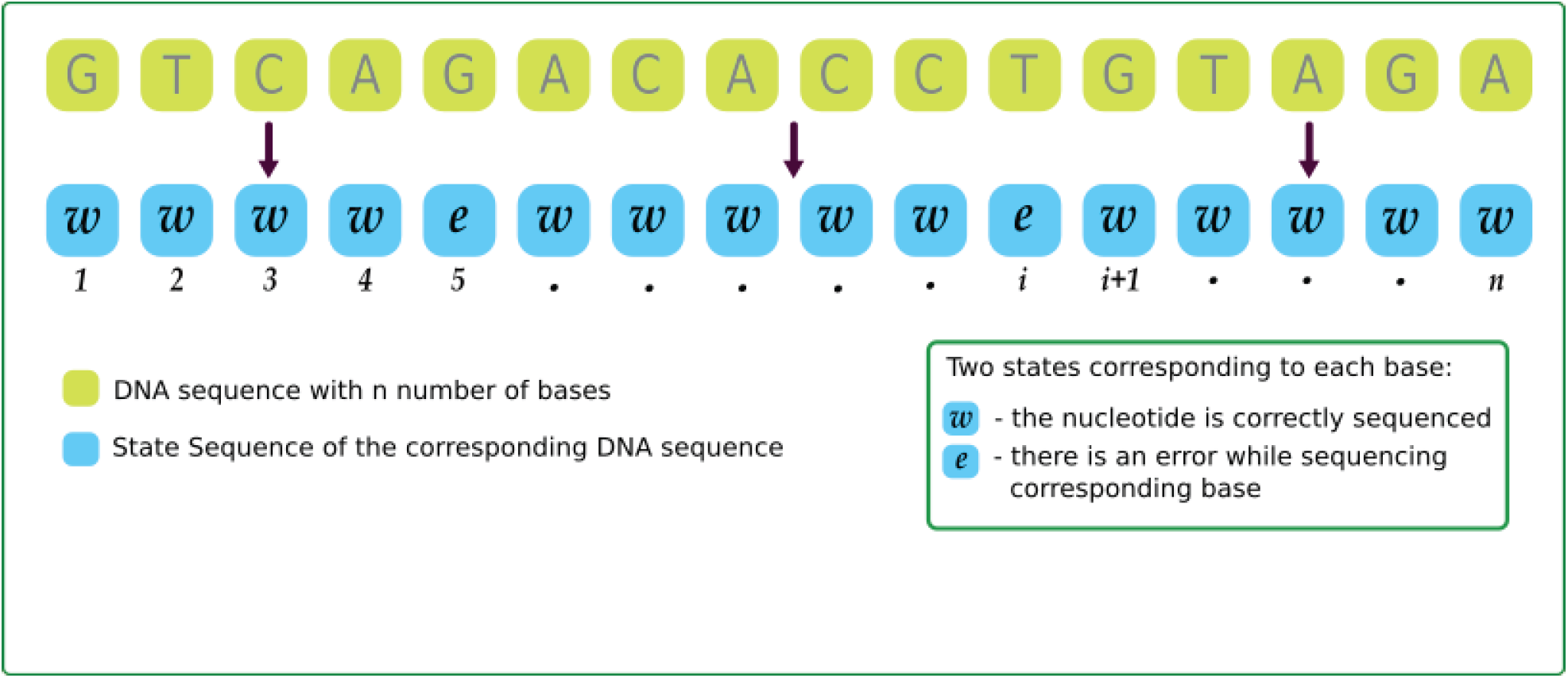
DNA sequence and its State Sequence where state is defined for each base in DNA sequence.Length of both the sequence is same

Now, if the probability of arrival of state ***e***(error at a position) is ‘*p*’ then the arrival of state ***w*** is ‘*1-p*’(the probability that base is correctly sequenced). The *p* value will be the average rate of error per nucleotide and therefore it has the same value for all the bases. Fox et al evaluated this error rate and reported as 0.1**[11]**. Since the error at every position is independent(as our assumption), it allows us to model error occurrence in the sequencing of DNA using Binomial Process with parameter *p*. If the length of the sequence is **n** and for finding the probability of total **k** errors in the sequence is given by:

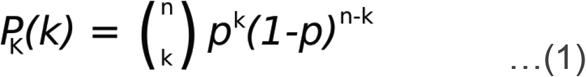

This is Binomial Distribution with parameter *p* as the probability of occurring error at a position and K is a random variable defined as the number of errors in the whole sequence of length **n**. In other words, it is the probability of **k** error states in the total state sequence of length **n**. For the complete sequence to be error less, K = 0 and the below equation gives the probability of the complete correct sequence.

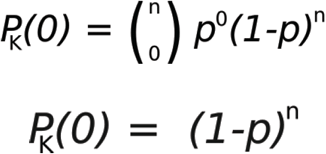

This is the ultimate probability of sequence to be accurate. As the length increases the overall probability decreases. This is because of (1-*p*)^*n*^. Error in the short length follows a binomial distribution (as shown in equation 1) and this would be our foundation of our whole mathematical model. We will extend it to find an accuracy of the genomic sequence and also studies based multiple sequences. Binomial Process can be approximated to the Poisson process for larger values of **n [7,8]**.

### POISSON PROCESS FOR GENOME SEQUENCE

Beyond Poisson approximation, we need to validate the properties of the poisson process with the occurrence of error in the long genome sequence.

1. Small interval probability - At a very small scale, the sequence is discrete where there are only two states ‘w’ and ‘e’. No two or more errors can occur at the same position. In addition to that, there is no feasibility of fractional error at a position. Therefore the probability of small interval can be given as:

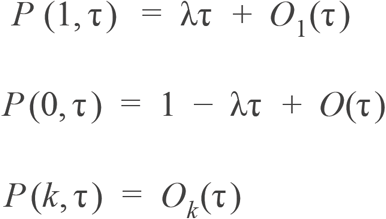 τ is the smallest interval in which the probability of one error occurrence is dependent on λ (the poisson rate). The probability of 2 errors is negligible in that interval τ.
2. Independence - The error arrival is independent of everything as our assumption.
3. Homogeneity - The probability of k arrivals will remain the same for all intervals of same length throughout the sequence. If the length τ _1_ and τ _2_ are same i.e. the number of bases in τ _1_ and τ _2_ are the same and therefore, the probability of *k* arrivals will remain the same for both the intervals**[7]**.

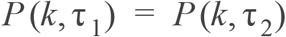

The probability distribution function can be given as:

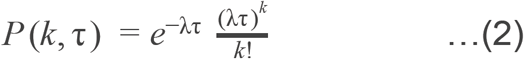

The equation (2) gives the probability of the *k* errors in a long DNA sequence (for e.g. Genomic sequence) of length τ. For the genome to be 100% accurate, the *k* must be 0. The accuracy(*k* = 0) decreases with the length as per equation (2).

### ERROR MODELLING IN FURTHER APPLICATION

Studies like GWAS, Genome annotation and cancer studies involve multiple sequences and their accuracy can be modeled by further extending equation (1) and equation (2) as follows:

1. After sequencing we need to compare it with other sequences. Therefore, alignment is one of the most fundamental parts of Computational Biology. The alignment efficiency is dependent on the algorithms as well as the sequencing quality. There are algorithms (for e.g. dynamic programming, BLAST) which are very efficient but error in sequencing can lead to variation in alignment. If the sequencing is 100 percent accurate, then the alignment is optimum and will produce proper results. So using the basis of errors in sequencing, we can calculate the probability of alignment to be fully accurate.
  a. For Shorter sequences: Error in alignment follows Binomial distribution by merging of two binomial processes. Consider an alignment involving two sequences. The error rate per nucleotide of the two sequences are **p** and **q** (Figure 3). The state sequence of both the sequence is shown in Figure 3. Using these two sequences, we are defining an *Alignment State Sequence* where two states are defined for every nucleotide position as ***w*** - when the alignment at that position is correct and *e* - when there is an error in alignment of respective nucleotide because of error in sequencing in any of the sequence. If there is an error in one sequence (at position *i*) then it will cause an alignment error for the same position.

**Figure 2:**
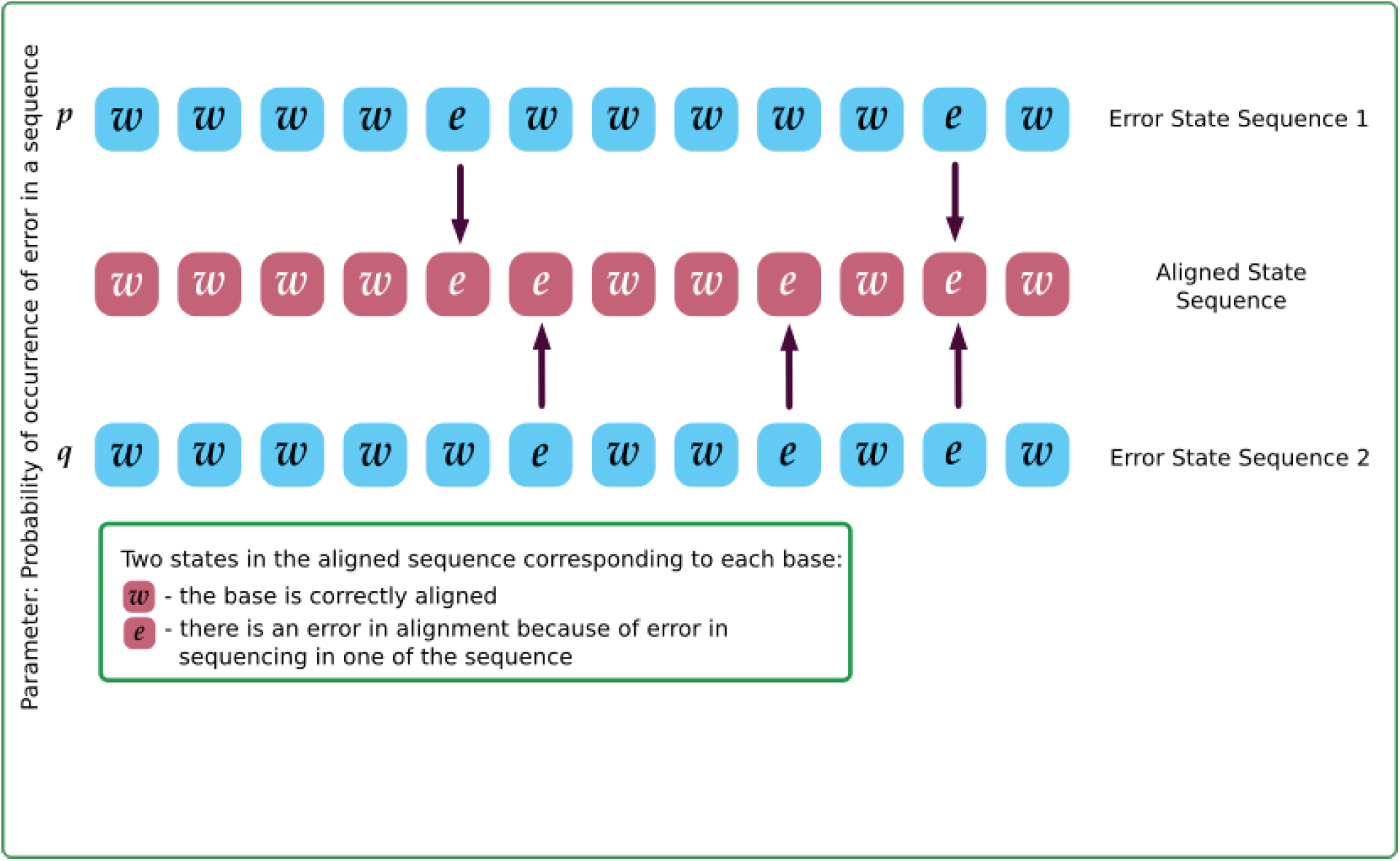
Two State Sequence defined for DNA sequences used in alignment where state is defined for each aligned base. Error in alignment follows Binomial distribution by merging of two binomial processes. This is an application of merging of two binomial processes. Now the probability of alignment to be fully correct is given by:

**Figure 3:**
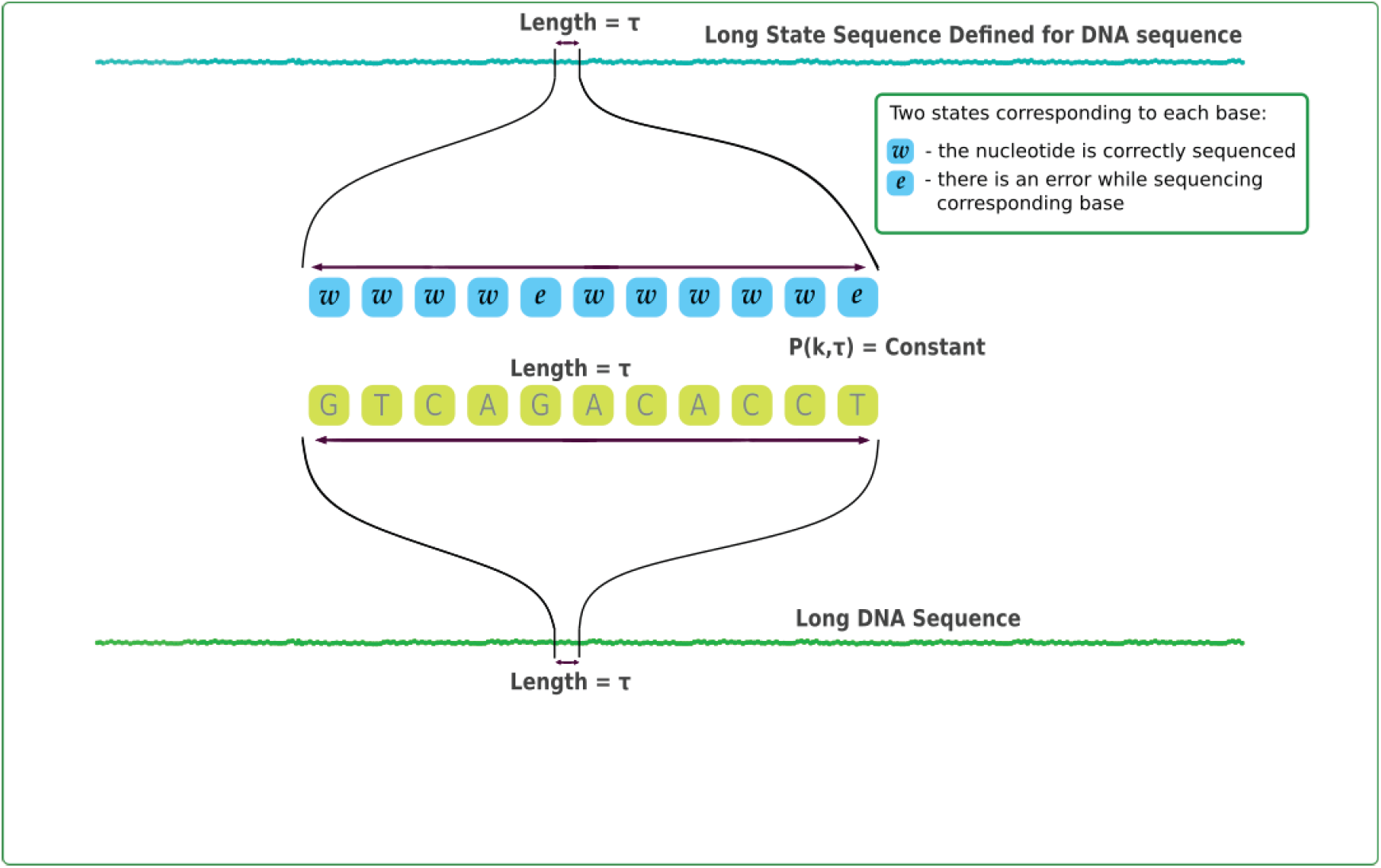
A long DNA sequence and its State Sequence. Each state is defined for a corresponding base at same position. The small length τ is a enlarged picture of both the sequences. (*P*_*w*_) Probability of w: P(not **e** in sequence 1)*P(not **e** in sequence 2) (*P* _*e*_) Probability of e: 1 - (*P*_*w*_)

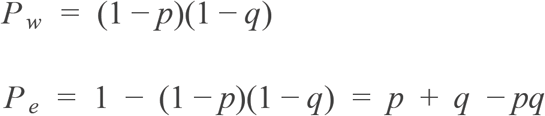 The result is also a binomial distribution but with different parameters as *P*_*e*_. Merged Binomial Distribution given as:

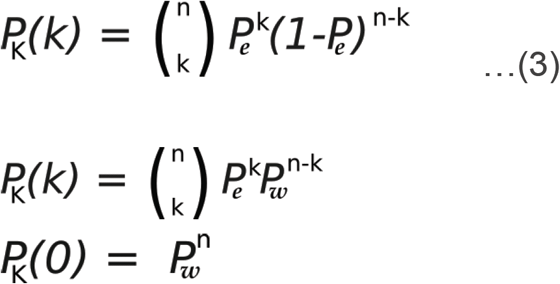 Special case when both the sequence are sequenced using same sequencer or same sequencing technology, then the parameter doubles as:

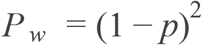
  b. GENOME ALIGNMENT: When two genomes are aligned, the resultant alignment is also a poisson process. Sequencing error in one sequence can cause an error in genome alignment. It can also be seen as a Poisson approximation of equation (3) for larger values of **n**. The accuracy of the genome alignment is a merged poisson process where the parameter will be as:

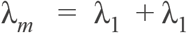 Where, λ_1_ and λ_2_ are the poisson parameters of the genome sequences and λ_*m*_ is the new poisson parameter for the error in aligned sequence aligned sequence. Special case: When both the sequences are sequenced using same technology or same sequencer, then their poisson parameter will remain same and equation (2) will become,

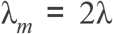

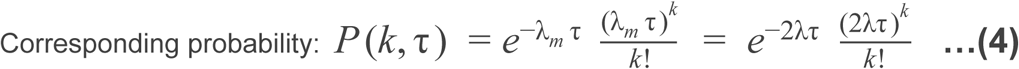
2. MOTIF DISCOVERY: Motif discovery involves multiple sequence alignment but of smaller length. Accuracy of a motif discovered can be calculated by using our basic Binomial Distribution model of errors in short sequence. It will again follow a Binomial Distribution by merging of multiple Binomial Processes. Considering a motif discovered using six DNA sequences (Figure 4). We defined a *Motif State Sequence* in a similar way with Alignment State Sequence as **c** - accurate motif base at a position and **e** - if the motif discovered is errorfull at that position because of error in sequencing in one of the sequences. Any error in one sequence would lead to inaccuracy in Position Weight Matrix and therefore, for the Motif discovered to be 100 percent accurate, all sequences to be accurate at every position.

*P*_*w*_ is the probability of ***w*** state at a position in Motif State Sequence i.e. Probability that the Motif base discovered is accurate at that position and *P*_*e*_ is the probability of state ***e*** at a position when there is an error and it is not accurate because of error in any sequence for that position.

**Figure 4:**
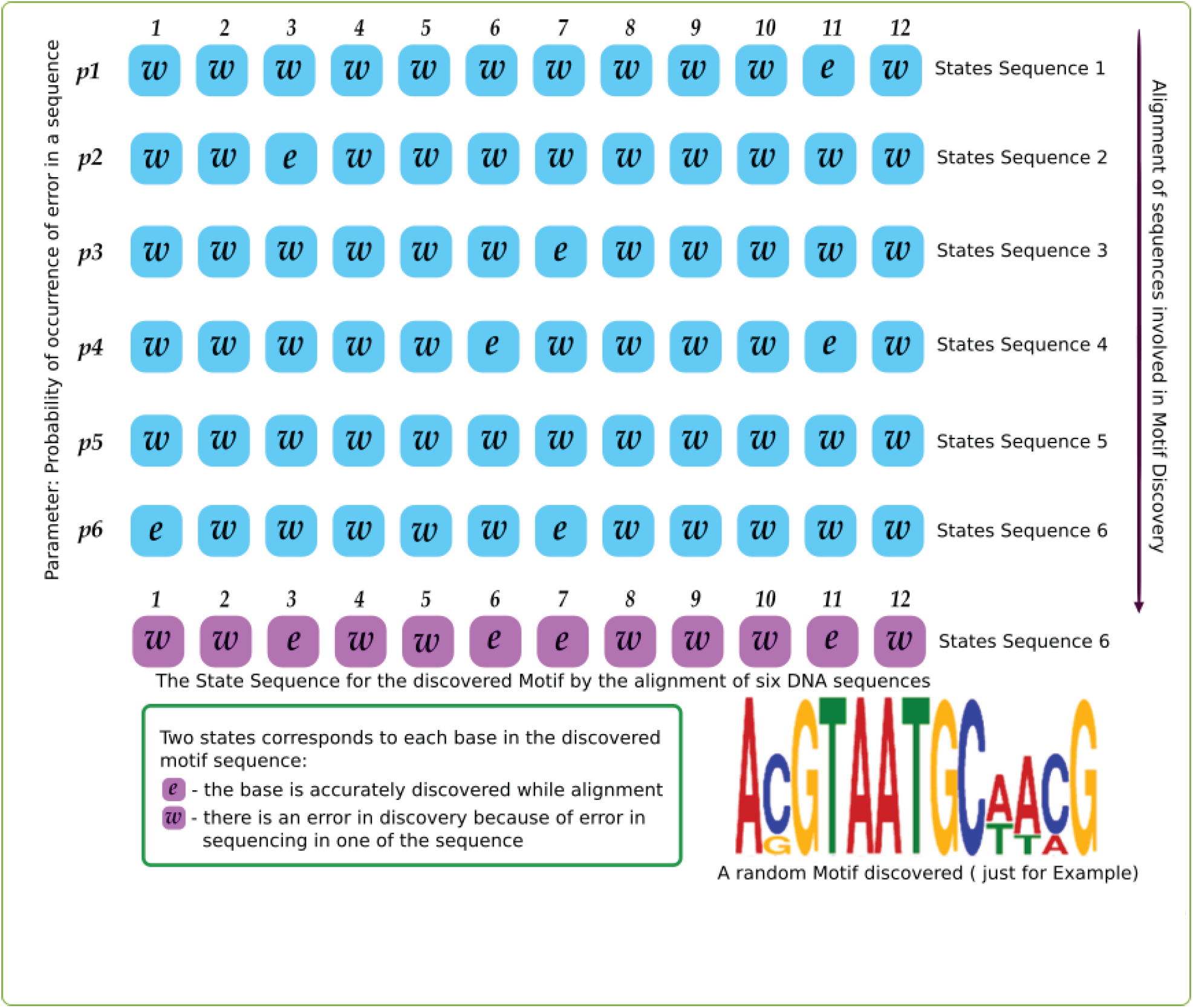
Six Sequences are aligned for finding a Motif. Above six sequences represents the Error State Sequence of the six DNA sequences. The last sequence is the Motif State Sequence for the discovered Motif.

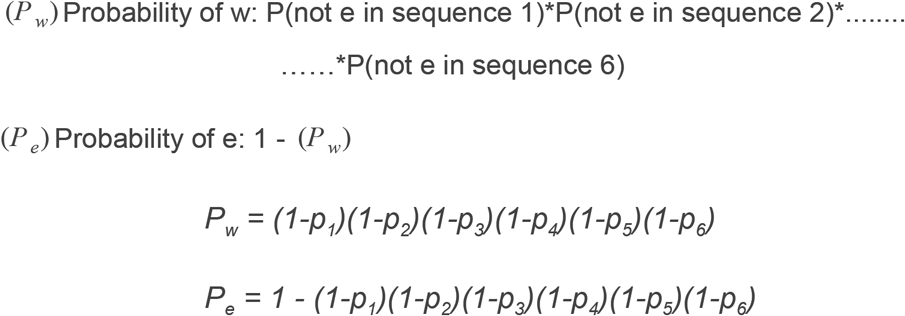

The Motif State Sequence will follow Binomial Distribution with parameter *P*_*w*_. It is similar to merging of two binomial processes in the alignment case. For generalised case when there are ***m*** sequences used for motif discovery:

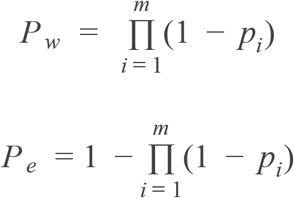

Special case when all the sequences are sequenced from same sequencer:

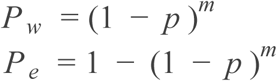

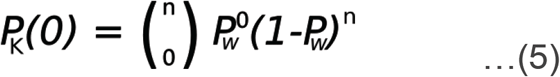

This is the Binomial Process for Motif Sate Sequence so that the motif sequence to be fully accurate. Equation (5) gives the probability of zero errors in motif. This idea of merging distribution can be applied to any study which involves alignment of multiple sequences and the Binomial Distribution can be further approximated to Poisson Distribution for longer sequences. Studies like GWAS and PANCancer compare massive genomic data for new findings in Disease Biology and the idea of merging can be used for systematically find the accuracy probability.

## Conclusion and Further Discussion

We presented a simple and very basic example of Probability in Biology. Errors in sequencing i.e. misreads follow a binomial distribution and it further can be approximated to the Poisson process for longer sequences. However, the insertion and deletion rates are 1000 times lower than substitution error rates and therefore less significant**[3]**. The model assumes that error arrival at a position is not dependent on error at other positions. Furthermore, errors in sequences can cause error in studies which are based on multiple sequences and they also follow Binomial – Poisson Distribution (for example – Alignment is a merging of two Binomial processes for short sequences and it further can be approximated to Poisson for long sequences (for example genomic sequence). It provides a systematic way to evaluate the accuracy in sequencing-based applications. Many error suppressing algorithms or techniques are there, and our Binomial Poisson model can provide further systematic understanding of error behavior in sequences so that more techniques for error removal can be developed with much effective suppression rates. The whole paper is focused on the basic mathematical biology and how very fundamental processes in probability can be applied to biology. There are further research possibilities for deeper understanding of errors in sequencing and its based application.

## Aknowledgement

We are thankful to edx.org and MIT-OCW for their online courses of Biology (MITx 7.00x) and Probability(MITx 6.041 and 6.012).

